# The role of β-defensin 103 (*DEFB103*) copy number variation in bull fertility

**DOI:** 10.1101/2024.08.27.609910

**Authors:** Ozge Sidekli, Edward J. Hollox, Sean Fair, Kieran G. Meade

## Abstract

Pregnancy rates for elite bulls used in artificial insemination (AI) can vary significantly and therefore the identification of molecular markers for fertility and targets to improve bull selection is important. β-defensins are peptides which have diverse regulatory roles in sperm function across multiple species. In this study, Holstein-Friesian bulls were screened based on field fertility data to identify two groups (High and Low fertility (HF and LF, respectively)) of n=10 bulls per group which were genotyped for copy number variation (CNV) in the *DEFB103* gene. Overall, low *DEFB103* copy number (CN) was associated with increased sperm motility across all bulls (n=20, p<0.05). As genetic diversity of *DEFB103* CN was only apparent in the LF group, three bulls per CNV class (low, intermediate and high CN) were chosen for functional analysis. Sperm from LF bulls with low CN exhibited higher binding to the oviduct epithelium *in vitro*, while high CN affected sperm membrane fluidity in non-capacitating conditions *in vitro* (p<0.05). To investigate the functional effect of *DEFB103* CNV on the uterine response *in vivo*, 18 heifers were inseminated with sperm from bulls with low, intermediate and high CN. Transcriptomic analysis on uterine tissue harvested 12 h post-insemination showed significant differential expression of 58 genes (FDR<0.1) involved in sperm migration, immune signalling and chemotaxis. These novel results confirm an important role for *DEFB103* CN in both sperm function and the uterine response to bull sperm, thereby potentially influencing pregnancy outcomes in cattle.

**Summary Sentence:** *DEFB103* copy number (CN) is associated with sperm motility and binding to the oviduct epithelium and uterine gene expression, thereby potentially influencing fertility outcomes.

## Introduction

Fertility is critically important for the livestock industry, as it directly affects production outcomes (Hales, 2019) as well as underpinning the economic sustainability of farming systems. This is particularly true within a seasonal calving system, where a cow must produce a calf per year and calve early in the spring to maximise the amount of milk produced from low-cost grass forage. Historically, intense genetic selection for production traits has resulted in a reduction in fertility [2], and although selection traits have broadened in recent years, sub-optimal fertility remains one of the principal reasons for premature culling of cows [3]. However, the role of the bull in determining successful pregnancy outcomes is often overlooked. Despite extensive *in vitro* quality controls checks of semen prior to its use in artificial insemination (AI), individual bull’s pregnancy rates can differ by up to 30% [4]. Various analytical techniques, such as computer assisted sperm analysis (CASA) and flow cytometry, have been used to elucidate variation in semen quality, with consequences for bull fertility. With these methods, although 47% of the variation in bull fertility in the field can now be explained by *in vitro* evaluations [5,6], a large component of bull fertility remains unaccounted for. Given that a single bull used in AI can inseminate thousands of females per season, it is clear that optimal fertility of the sire is of critical importance to herd fertility outcomes [4].

The reasons for differences in fertility between AI bulls are likely to be multifactorial, involving issues with sperm transport in the female reproductive tract, fertilization, pre-implantation embryonic development and conceptus elongation as well as development of the embryo and placenta after recognition of pregnancy [7]. Understanding the molecular mechanisms involved in sperm maturation, and also between sperm and the female reproductive tract underpinning sperm-oviduct and uterine interactions, will involve identifying proteins which determine fertility outcomes. These proteins may improve diagnosis of infertility as well as identify targets for enhanced breeding in livestock.

Sperm are coated with many different multifunctional peptides, including β-defensins. β-defensins are peptides defined by a conserved six-cysteine motif, which forms three disulphide bridges in a 1-5, 2-4, 3-6 arrangement [8]. Some β-defensins are short – typically ∼60 amino acids in length, while others have an extended C-terminal tail, containing potential O- and N-glycosylation sites [9,10]. Initially characterised as antimicrobial peptides, β-defensins also have important roles as inflammatory mediators and in male reproductive traits. For example, mice with deletions of β-defensin genes show male infertility [11], and a variant of β-defensin 126 is associated with infertility and reduced sperm motility in men [12,13]. The functional role of β-defensin 126 has also been explored in macaques, where it is secreted in the epididymis, and binds to sperm with the heavily-glycosylated C-terminal tail which forms part of the negatively-charged sperm glycocalyx [14]. It facilitates penetration of cervical mucus and mediates attachment of the sperm to oviduct epithelia [14–16]. In cattle, β-defensin 126 is adsorbed on the tail and post-acrosomal region of the sperm and promotes sperm motility [17]. In addition, β-defensins may also impact on the regulation of the immune response in the female reproductive tract to improve sperm survival and/or pregnancy outcomes. A study directly comparing the *in vivo* uterine response of sperm from bulls with different fertility phenotypes showed an upregulation of genes associated with pregnancy establishment in response to sperm from the high fertility bulls [18], although the role of β-defensins was not assessed.

The β-defensin gene family are known to show recent duplications and genetic divergence, forming clusters of often-closely related genes. Several mammalian lineages show evidence of extensive copy number variation (CNV) of β-defensin genes between individuals [19]. This observation includes cattle [20,21], where *DEFB103*, in particular, shows extensive multiallelic CNV and shows a diploid copy number (CN) range of 1 to ∼29 across seven different cattle breeds [21]. Studies of β-defensin genes in cattle [22] and rats [23] have shown regional expression patterns of β-defensins in the epididymis. In rats, *Defb14*, an ortholog of *DEFB103*, is expressed in both the testis and cauda epididymis [23]. We have found that *DEFB103*, which encodes BBD103, is expressed in the caput of the epididymis in cattle; is upregulated in testes during sexual maturation from calves to bulls and that mRNA expression levels negatively correlate with gene CN [21]. Taken together, we hypothesize that *DEFB103* CNV is a strong genetic candidate for affecting bull fertility, with the encoded BBD103 protein potentially influencing fertility outcomes and related phenotypes.

## Methods

### Ethical approval

The procedures were established in compliance with the Cruelty to Animals Act (Ireland 1876, as modified by European Communities regulations 2002 and 2005) and the European Community Directive 86/609/EC.

### Reagents

Unless specified, all chemical substances were obtained from Merck, Arklow, Co Wicklow, Ireland.

### Bull selection

Data on the field fertility of a population of Holstein-Friesian bulls (n = 840) used in Ireland were obtained from the Irish Cattle Breeding Federation (ICBF) database. Each bull had a minimum of 500 inseminations based on an adjusted sire fertility model (Berry et al.,2011). Adjusted bull fertility was defined as pregnancy to a given service identified retrospectively either from a calving event or where a repeat service (or a pregnancy scan) deemed the animal not to be pregnant to the said service. Cows and heifers that were culled or died on farm were omitted. These raw data were then adjusted for factors including semen type (frozen, fresh), cow parity, days in milk, month of service, day of the week when serviced, service number, cow genotype, herd, AI technician, and bull breed and were weighted for number of service records resulting in an adjusted pregnancy rate centered at 0%. For this study, bulls classified as having HF had an average adjusted fertility score of +6.4 whereas, those classified as LF was -6.5. Genotyping data of bulls for *DEFB103* CNV was previously obtained [21] using digital droplet PCR (ddPCR). Among 20 Holstein Friesian bulls, 9 LF sires were categorized based on their *DEFB103* CNV genotype (into three groups: low CN (n=3 bulls), intermediate CN (n=3 bulls), and high CN (n=3 bulls). In the *DEFB103* genotype, which we previously defined as multiallelic, a diploid CN of 6 was used as the reference, and distinct threshold values of <4.5 and >7.5 were used to define low CN and high CN of this gene, respectively. Among LF bulls with low, intermediate and high CNV, mean adjusted pregnancy rates were -8.96%, -3.96% and -7.8%, respectively.

Data were provided by Stiavnicka et al., (2023) to examine the effect of *DEFB103* CNV on various sperm motility and kinematic parameters (assessed using CASA), as well as functional traits including viability, membrane fluidity, and acrosomal status (assessed using flow cytometry) in vitro under both capacitating and non-capacitating conditions. modified Tyrode medium (mTALP) for capacitation conditions, which contains key ingredients such as NaHCO3, CaCl2, BSA and Heparin, and non-capacitation medium (NCM), which is similar to but lacks these ingredients, containing increased NaCl and polyvinyl alcohol. They washed and resuspended frozen-thawed sperm samples in either NCM or mTALP and incubated them under conditions specific to capacitation or non-capacitation. They assessed sperm viability, membrane fluidity, and acrosome integrity using flow cytometry, induced hyperactivation with caffeine, and evaluated it using CASA. Additionally, they triggered the acrosome reaction with Calcium Ionophore A23187 and a vehicle control with dimethyl sulfoxide, and assessed it post-incubation.

### Ability of sperm from bulls with different *DEFB103* CNV genotypes to swim through artificial mucus

This straw was subjected to washing and subsequent centrifugation at 300 *g* for 5 minutes. The concentration of sperm was adjusted to 10 x 10^6^ per mL. The ability of sperm to penetrate cervicovaginal mucus was assessed using the artificial mucus method previously described by Al Naib et al. (2011). Briefly, artificial mucus was formulated by diluting a sodium hyaluronate solution (MAP-5, Lab-stock MicroServices, Ireland) with phosphate-buffered saline, resulting in a final concentration of 6 mg of sodium hyaluronate per mL. Flattened capillary tubes (measuring 0.3 mm x 3.0 mm x 100 mm; Composite Metal Services Ltd, UK) were marked at 10 mm intervals from 10 to 90 mm. These tubes were subsequently filled with artificial mucus and sealed at one end. Two vertically positioned capillary tubes were introduced into a 1.5 mL eppendorf tube, each containing 100 µL of hoeschst stained sperm from one bull at a concentration of 10 x 10^6^ sperm per mL. Thus, for each replicate, each bull sperm was represented by two capillary tubes. The tubes were incubated at 37°C for 1 h, then placed on a hotplate at 45°C for 1 min to immobilize the sperm. Sperm were counted across the width of the tube within one field of view wide, at 10 mm intervals using a fluorescent microscope (10x; Olympus BX 60).

### Ability of sperm from bulls with different *DEFB103* CNV genotypes to bind to oviductal explants *in vitro*

The *in vitro* oviductal epithelium binding capacity of sperm obtained from LF bulls, characterized by the *DEFB103* CNV genotype (low CN (n=3), intermediate CN (n=3) and high CN (n=3)) was performed as per the method outlined by Lyons et al. (2018). Each bull’s thawed semen was diluted with a prepared stock culture containing medium 199 (M199) supplemented with fetal bovine serum (10%) and gentamicin sulfate (0.25 mg/mL). The sperm concentration was adjusted to 5 x 10^6^ sperm per mL. Reproductive tracts were collected from non-pregnant heifers (n = 9) at a commercial abattoir. Various stages of the oestrous cycle were represented, as previous literature has indicated that the stage of the oestrous cycle does not impact sperm binding *in vitro* [24]. Tissue explants were taken from the isthmic portion of the oviduct, and the explants from both oviducts were combined for each individual tract. Explants were prepared by cleaning oviducts, isolating the isthmic segment, and releasing epithelial cells through gentle compression and fragmentation. These were cultured in M199 medium supplemented with fetal calf serum and gentamicin sulfate, incubated to form everted vesicles, and used within 5 h of slaughter. For the binding assay, explants were combined with sperm (5 X 10^6^ sperm per mL) pre-stained with Hoechst 33342 dye for enhanced binding visualisation [16]. After incubation and removal of loosely bound sperm by gently pipetting through two 75 μL droplets of M199 media, samples were examined under a fluorescent microscope. A total of ten explants from each treatment were randomly evaluated, and the density of sperm binding determined by calculating the number of bound spermatozoa per 0.1 mm^2^ of explant surface.

### *In vivo* sperm migration into the oviduct and uterine transcriptomic response in bulls with different *DEFB103* CNV genotypes

#### Heifer selection

Eighteen 24- to 30-month-old cross-bred heifers (Charolais, Limousin, Aberdeen Angus and Simmental) had their oestrus cycles synchronized and were inseminated using a fixed-time AI protocol. The heifers were blocked by breed and age into three distinct treatment groups and inseminated with semen from 3 bulls from each treatment, namely, low CN (7 heifers), intermediate CN (6 heifers), and high CN (5 heifers). The synchronization protocol entailed a seven-day intravaginal progesterone device (CIDR®), accompanied by intramuscular administration of gonadotrophin-releasing hormone (Ovarelin; 2 mL) at CIDR® insertion. Prostaglandin F2 alpha (Enzaprost; 5 ml) was administered intramuscularly to induce luteolysis 24 h before CIDR removal. At 54 (+/- 1) h post-CIDR® removal, heifers received a single fixed-time insemination of frozen–thawed semen. Ovarelin (2 mL) was administered intramuscularly at the time of AI.

#### Tissue collection

Twelve hours post insemination, heifers were slaughtered at a commercial abattoir. Reproductive tracts were promptly collected post-mortem, and the uterine horn on the side of the ovulated ovary was longitudinally opened with sterile scissors. Tissue samples were acquired from the inter-caruncular region of the uterine horn near the uterine body using a sterile 8-mm biopsy punch. The endometrium was then dissected away from the myometrium. Endometrium samples were flash frozen in liquid nitrogen, transported to the laboratory, and stored at −80°C.

Total RNA isolation, RNA library preparation and sequencing was conducted by Biomarker Technologies (BMKgene, Munster, Germany). Paired-end (150bp) sequencing was performed on an Illumina Novaseq 6000 sequencer to a raw read depth of approximately 50 million total reads (303.20 Gb) per sample. Percentage of Q30 bases in each sample was above 98.18 %.

#### Quality control, mapping, and differential read count quantification

Raw sequence reads were obtained in FASTQ format and their quality was evaluated using FastQC (v.0.12.1). The sequences from all samples were then quality-trimmed and cleared of adaptor sequences using the BBDuk Java package. For alignment, trimmed reads were semi-mapped to the bovine reference genome (ARS-UCD1.2) using Salmon [25], and uniquely mapped read counts per Ensembl (version 104) annotated gene/transcript were estimated using the Salmon–quant mode option. To ensure robustness, only genes with a minimum of five reads across all samples were considered in subsequent analyses. MultiQC analysis was then conducted to derive essential statistics on the percentage of reads counted in the transcriptome. The Tximport function (v.1.3.9) [26] in R 4.3.1 summarized the results at the per-transcript gene level.

Differential gene expression analysis, along with data transformations and visualization, was performed using DeSeq2 (v3.18) [27] in R 4.3.1. Samples were clustered based on variance-stabilizing transformed data and visualized through principal component analysis (PCA). Differentially expressed gene (DEG) lists were created using a negative binomial generalized linear model, and comparisons were conducted among uterine biopsy groups. P values were adjusted for multiple comparisons using the Benjamini and Hochberg (B–H) method. Genes with an adjusted P-value (False Discovery Rate (FDR)) <0.1 were considered differentially expressed and used for further data exploration and pathway analysis.

Gene Ontology (GO) and Kyoto Encyclopedia of Genes and Genomes (KEGG) pathway analysis were used for all DEG regardless of specific *DEFB103* CN groups. GO term was analyzed using the Gene Ontology knowledge base [28,29]. In the pathway analysis, the analysis of DEGs was performed using ClusterProfiler (v3.18.0) based on the KEGG database [30]. In this analysis, GO terms and pathway analysis were defined as P adjusted (FDR) < 0.05.

### Recovery of sperm from bovine oviduct

Oviducts from the 18 heifers (total n = 36 paired oviducts) were collected for sperm recovery. One end of each oviduct was tied with a cotton thread, spTALP solution with heparin (10 μg/mL) was prepared and injected into the oviducts, and the oviducts were closed. After 60 minutes of incubation, the oviducts were milked, and sperm concentration was assessed using a haemocytometer.

### Statistical analysis

Pearson correlation coefficients were calculated for the relationship between *DEFB103* CNV with CASA and Flow Cytometry data, and a linear regression analysis was performed between the results via GraphPad Prism (v 9.5.1). All other data were checked for normality of distribution and analyzed by univariate analysis of variance (ANOVA) with Bonferroni post-hoc tests in GraphPad Prism (v 9.5.1). Statistical significance level was set at p <0.05. All results are presented as mean ± standard error of the mean.

## Results

### Impact of *DEFB103* CNV on sperm kinematics, mucus penetration, and oviductal epithelial binding

The objective of the study was to assess the impact of *DEFB103* CNV on sperm functional parameters, mucus penetration, and binding to the oviduct epithelium. Low *DEFB103* CN was strongly associated with increased total motility (p<0.05) and increased progressive motility (*p*<0.05). Low *DEFB103* CN was associated with lower straight-line velocity (VSL; *p*<0.05), average path velocity (VAP; *p*<0.05), and higher sperm-oviduct binding capacity (*p*<0.05; Table 2 and 3; Figure 1). Moreover, the percentage of viable sperm with high membrane fluidity in non-capacitated conditions demonstrated that high *DEFB103* CN was associated with higher membrane fluidity compared to the intermediate CN group (*p*< 0.05). There was no effect of *DEFB103* genotype on the ability of sperm to penetrate artificial mucus. Taken together, these findings suggest that low *DEFB103* CN is associated with increased sperm binding to the oviduct epithelium and improved motility, while high *DEFB103* CN tends to affect membrane fluidity under *in vitro* capacitating conditions.

**Figure 1:**
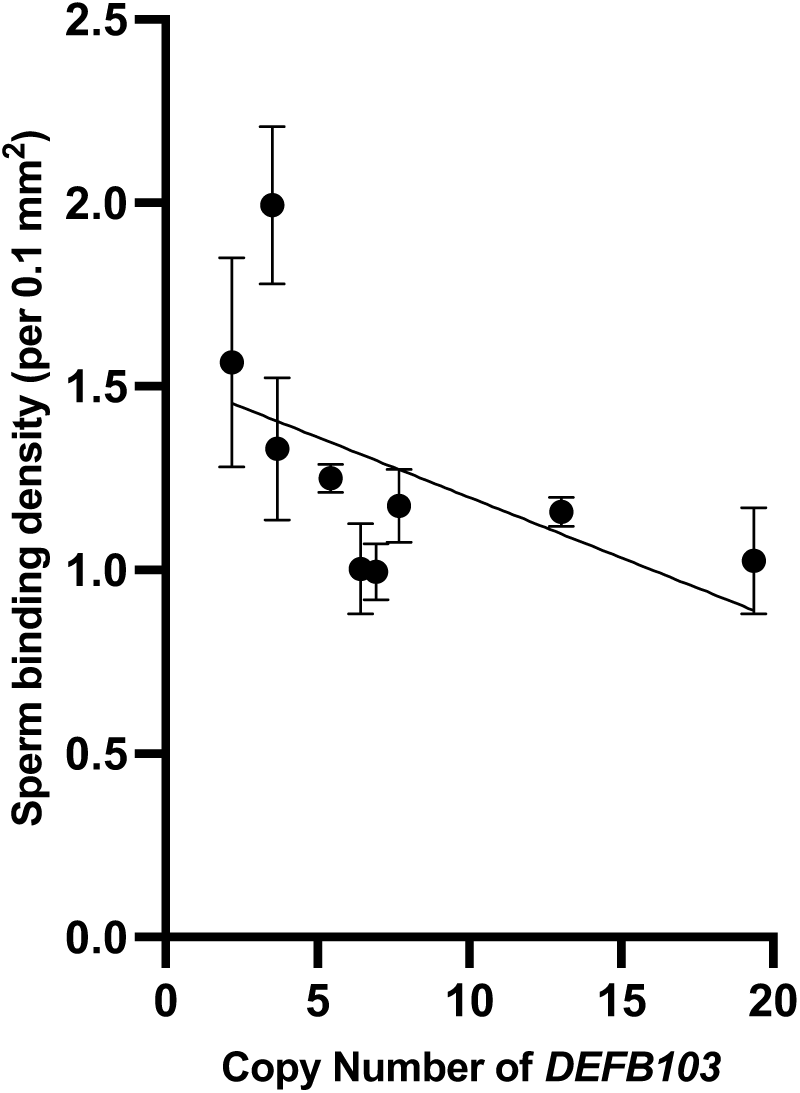
The ability of frozen-thawed sperm from low fertility Holstein-Friesian bulls with varying *DEFB103* copy number variation (CNV) to bind to the oviductal epithelial *in vitro.* Mean and standard errors of three replicates are shown for low fertility bulls (n=9). r=-0.5, p=0.006.

**Table 1:** Individual-specific data on copy number variation (CNV) for the *DEFB103* gene across all three loci (including *DEFB103A, DEFB103A-Like,* and *DEFB103B*) and fertility phenotypes namely, low fertility (LF) and high fertility (HF) in Holstein Friesian bulls (n=10 per group). Evaluating the combined loci of *DEFB103*, the classification of CNV for the diploid genome categorizes them as having low, intermediate, or high

| Individual bull ID | Fertility Status | Adjusted Pregnancy Rate (%) | Number of Inseminations which the Adjusted Pregnancy Rate is based on | *DEFB103* CN | *DEFB103* CNV status |
| --- | --- | --- | --- | --- | --- |
| H1 | HF | 6.8 | 100288 | 2.22 | Low |
| H2 | HF | 6.5 | 3912 | 3.04 | Low |
| H3 | HF | 5.8 | 11459 | 2.76 | Low |
| H4 | HF | 6.8 | 12424 | 5.82 | Intermediate |
| H5 | HF | 7.1 | 34973 | 2.34 | Low |
| H6 | HF | 6.2 | 17441 | 3.09 | Low |
| H7 | HF | 6.7 | 1041 | 2.94 | Low |
| H8 | HF | 6.2 | 5119 | 4.38 | Low |
| H9 | HF | 6 | 8637 | 2.13 | Low |
| H10 | HF | 6.7 | 37849 | 5.82 | Intermediate |
| L1 | LF | -8.5 | 1034 | 2.19 | Low |
| L2 | LF | -4.3 | 23811 | 6.42 | Intermediate |
| L3 | LF | -3 | 568 | 6.93 | Intermediate |
| L4 | LF | -9.1 | 740 | 3.51 | Low |
| L5 | LF | -4.6 | 597 | 5.43 | Intermediate |
| L6 | LF | -3.5 | 1195 | 4.35 | Low |
| L7 | LF | -9.3 | 519 | 3.69 | Low |
| L8 | LF | -12.3 | 1477 | 13.05 | High |
| L9 | LF | -7.3 | 1772 | 19.35 | High |
| L10 | LF | -3.9 | 980 | 7.68 | High |

**Table 2:** Correlation between *DEFB103* copy number variation (CNV) and sperm functional parameters in Holstein-Friesian bulls with high and low fertility phenotypes (n=10 per group). P values <0.05 are shown in bold.

| Parameter | Pearson r | P value |
| --- | --- | --- |
| **Computer Assisted Sperm Analysis (CASA)** |  |  |
| Total motility (%) | -0.64 | **0.003** |
| Progressive motility (%) | -0.54 | **0.015** |
| Curvilinear velocity (VCL; $\mu\text{m/s}$) | 0.07 | 0.76 |
| Straight-line velocity (VSL; $\mu\text{m/s}$) | 0.33 | 0.16 |
| Average path velocity (VAP; $\mu\text{m/s}$) | 0.29 | 0.21 |
| Hyperactivation (%) | -0.34 | 0.14 |
| **Flow cytometry** |  |  |
| Viable (%) in capacitating conditions | -0.29 | 0.21 |
| Acrosome reacted (%) in capacitating conditions | -0.21 | 0.38 |
| Acrosome intact (%) in capacitating conditions | 0.20 | 0.38 |
| High membrane fluidity (%) in capacitating conditions | -0.13 | 0.57 |
| Viable (%) in non-capacitating conditions | -0.18 | 0.44 |
| Viable, acrosome react (%) in non-capacitating conditions | 0.01 | 0.94 |
| Viable, acrosome intact (%) in non-capacitating conditions | -0.01 | 0.94 |
| Viable, high membrane fluidity (%) in noncapacitating conditions | -0.06 | 0.78 |

### Effect of *DEFB103* copy number variation on sperm migration to the oviducts and uterine transcriptomic response

Given the effect of low CNV on sperm phenotype, in particular on sperm oviductal binding, we investigated the effect of the sperm from bulls with different *DEFB103* CN on sperm migration to the oviducts and uterine response *in vivo*. We confirmed that our results are not confounded by differing efficiencies of sperm reaching the ipsilateral and collateral uterine horns of heifers after AI, as there was no relationship between sperm concentration recovered by flushing and the *DEFB103* CN of the bull (Figure 2).

**Figure 2.**
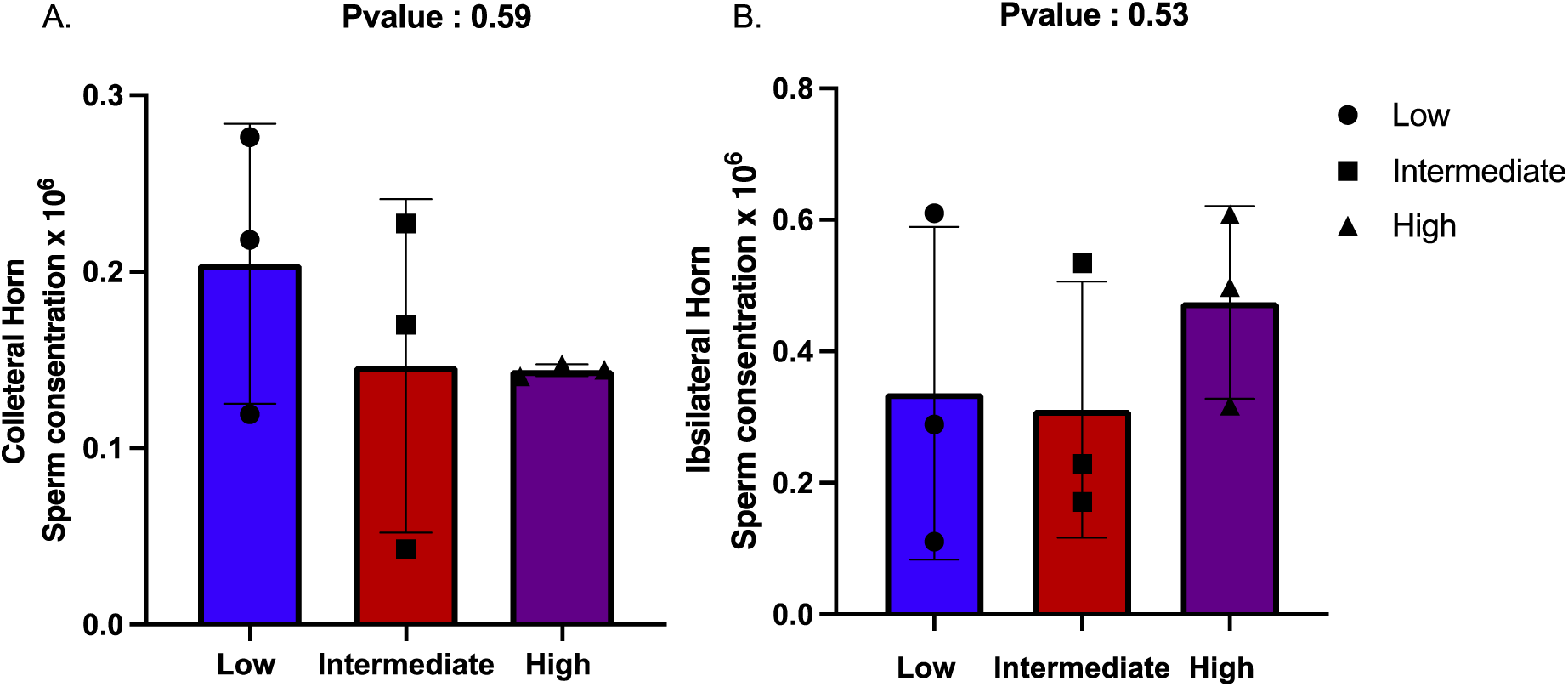
The ability of frozen-thawed sperm from low fertility Holstein-Friesian bulls (n=3 per group) with varying *DEFB103* copy number to populate the oviduct *in vivo*. (A) Sperm recovered from the contralateral horn, and (B) Sperm recovered from the ipsilateral horn after flushing. The X-axis represents the genotype, while the Y-axis indicates the sperm concentration (x 10^6^).

Principal component analysis of global transcript levels from the 18 different heifers uterus tissue showed no clear clustering based on the *DEFB103* CN of the bull whose semen was used for insemination (data not shown) indicating an absence of global differences in the transcriptome due to sperm treatment. This is likely accounted for by the complexity of diverse sperm-uterine interactions occurring simultaneously as well as the heterogeneity due to the use of different heifers. To identify more subtle differences between treatments, three pairwise comparisons were performed: 1. between heifers inseminated with semen from high *DEFB103* CN bulls and those inseminated by low CN bulls; 2. between heifers inseminated with semen from high *DEFB103* CN bulls and those inseminated by intermediate CN bulls; and 3. between heifers inseminated with semen from intermediate *DEFB103* CN bulls and those inseminated by low CN bulls.

A total of 58 genes were differentially expressed in the uterus post-insemination using sperm from *DEFB103* bulls using a threshold of FDR<0.1 (Full list shown in Supplementary Table S1). Some of these genes had a low fold change and those with a FC<-2 or FC > 2 are shown in Table 4. In comparison 1, the high CN group showed increased expression of 5 genes (*KYAT3, CADM3, SEMA3C, SEMA3A*, and *SLC35F1*) compared to the low CN group, while the expression of 6 genes (*SKAP2, RPS3A, TGIF1*, *EFR3B,* and *CYP2C87*), including Calponin 3 (*CNN3*), was decreased in same comparison. Notably, *RPS3A*, which is an integral part of ribosomal proteins and critical for the protein synthesis process [31], and *CNN3*, which encodes an actin binder involved in actin cytoskeleton regulation, cell adhesion, and tissue remodelling [32], showed 256- and 512-fold decreases in expression levels, respectively. This reduction may hinder cell adhesion and reorganization, thus posing difficulties for embryo implantation and subsequent development.

**Table 3:** Effect of *DEFB103* copy number variation (CNV) on sperm functional parameters assessed using computer assisted sperm analysis (CASA), flow cytometry, mucus penetration and sperm binding to oviductal epithelial cells *in vitro* in low fertility Holstein-Friesian bulls. Values are mean ± s.e.m. n=3 bulls per group. P values <0.05 are shown in bold.

| Parameter | CNV status |  |  | Comparisons of each CN group (*P* value) |  |  |
| --- | --- | --- | --- | --- | --- | --- |
|  | Low | Intermediate | High | High vs Low | High vs Intermediate | Low vs Intermediate |
| **CASA** |  |  |  |  |  |  |
| Total motility (%) | 57.2±4.39 | 50.6±5.75 | 47.2±4.56 | 0.82 | >0.99 | >0.99 |
| Progressive motility (%) | 42.4±4.32 | 40.4±5.66 | 36.9±4.49 | >0.99 | >0.99 | >0.99 |
| Curvilinear velocity (VCL; mm/s) | 89.7±6.70 | 107.9±5.20 | 103.7±9.16 | 0.42 | >0.99 | 0.21 |
| Straight-line velocity (VSL; mm/s) | 58.9±4.98 | 80.7±5.02 | 79.0±9.49 | 0.11 | >0.99 | **0.04** |
| Average path velocity (VAP; mm/s) | 69.9±4.42 | 90.7±4.62 | 84.1±8.67 | 0.14 | 0.58 | **0.03** |
| Hyperactivation (%) | 6.8±3.31 | 3.7±1.11 | 4.5±2.39 | >0.99 | >0.99 | >0.99 |
| **Flow cytometer** |  |  |  |  |  |  |
| Viable (%) in capacitated condition | 14.2±3.90 | 10.3±3.86 | 14.4±3.02 | 0.6 | 0.99 | 0.57 |
| Acrosome reacted (%) in capacitated condition | 10.1±3.12 | 15.1±4.10 | 5.2±1.19 | 0.79 | 0.09 | 0.77 |
| Acrosome intact (%) in capacitated g condition | 89.8±3.12 | 84.8±4.10 | 94.8±1.19 | 0.79 | 0.09 | 0.77 |
| High membrane fluidity (%) in capacitated condition | 13.1±3.91 | 5.6±1.31 | 8.6±1.32 | 0.29 | 0.56 | 0.051 |
| Viable (%) in noncapacitated condition | 21.0±4.11 | 16.0±1.62 | 21.1±2.51 | >0.99 | 0.44 | 0.47 |
| Acrosome reacted (%) in noncapacitated condition | 1.7±0.82 | 2.8±1.32 | 1.0±0.11 | >0.99 | 0.49 | >0.99 |
| Acrosome intact (%) in noncapacitated condition | 98.2±0.85 | 97.1±1.33 | 98.9±0.11 | >0.99 | 0.49 | >0.99 |
| High membrane fluidity (%) in noncapacitated condition | 3.7±1.42 | 2.4±1.2 | 5.6±0.90 | 0.32 | **0.003** | 0.17 |
| **Mucus penetration** | 113.5±13 | 122.0±59 | 122.5±22.93 | 0.94 | 0.73 | 0.55 |
| **Sperm binding to oviductal epithelium** | 1.6±0.18 | 1.0±0.08 | 1.1±0.04 | 0.05 | 0.97 | **0.03** |

**Table 4.**
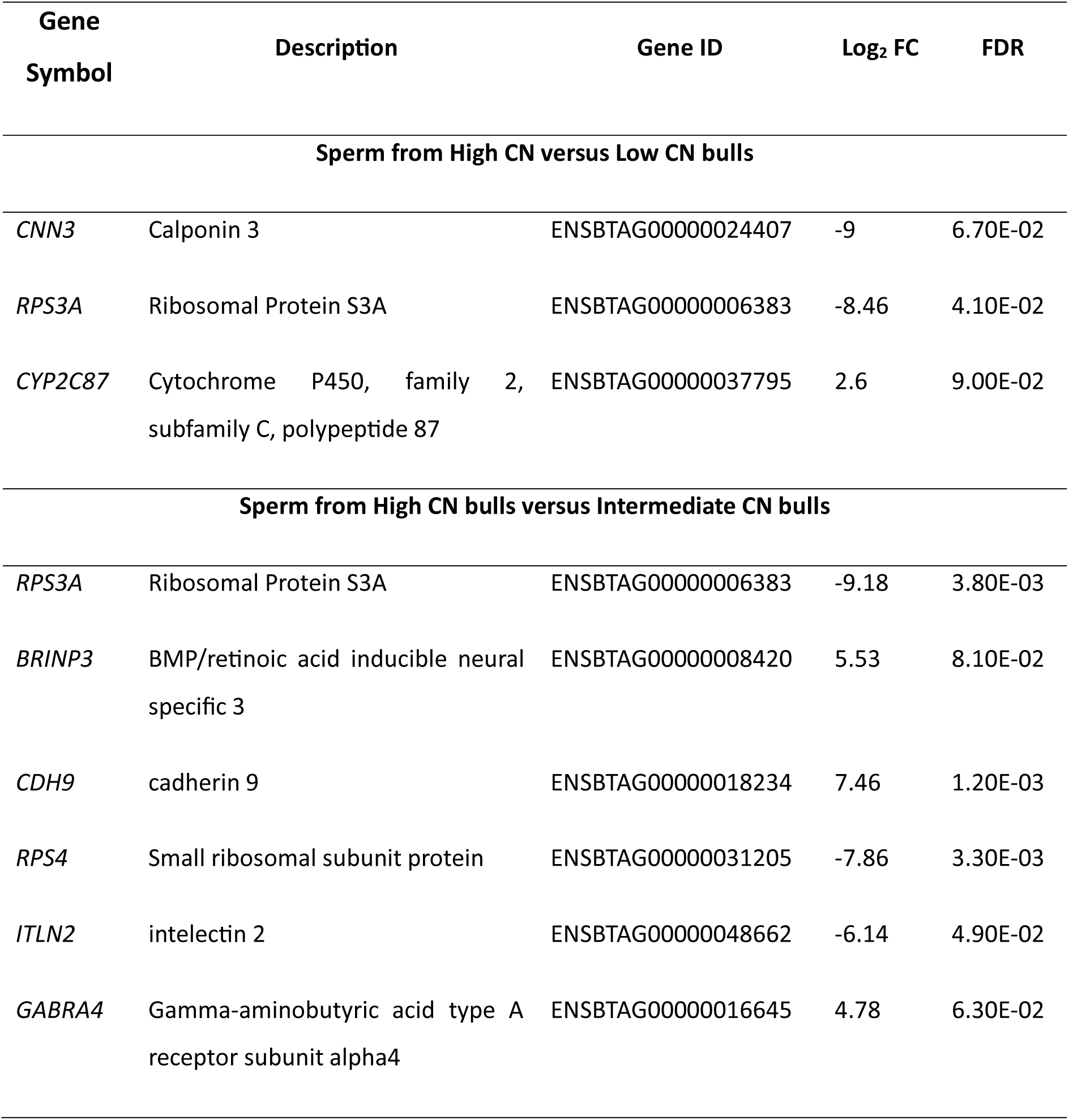
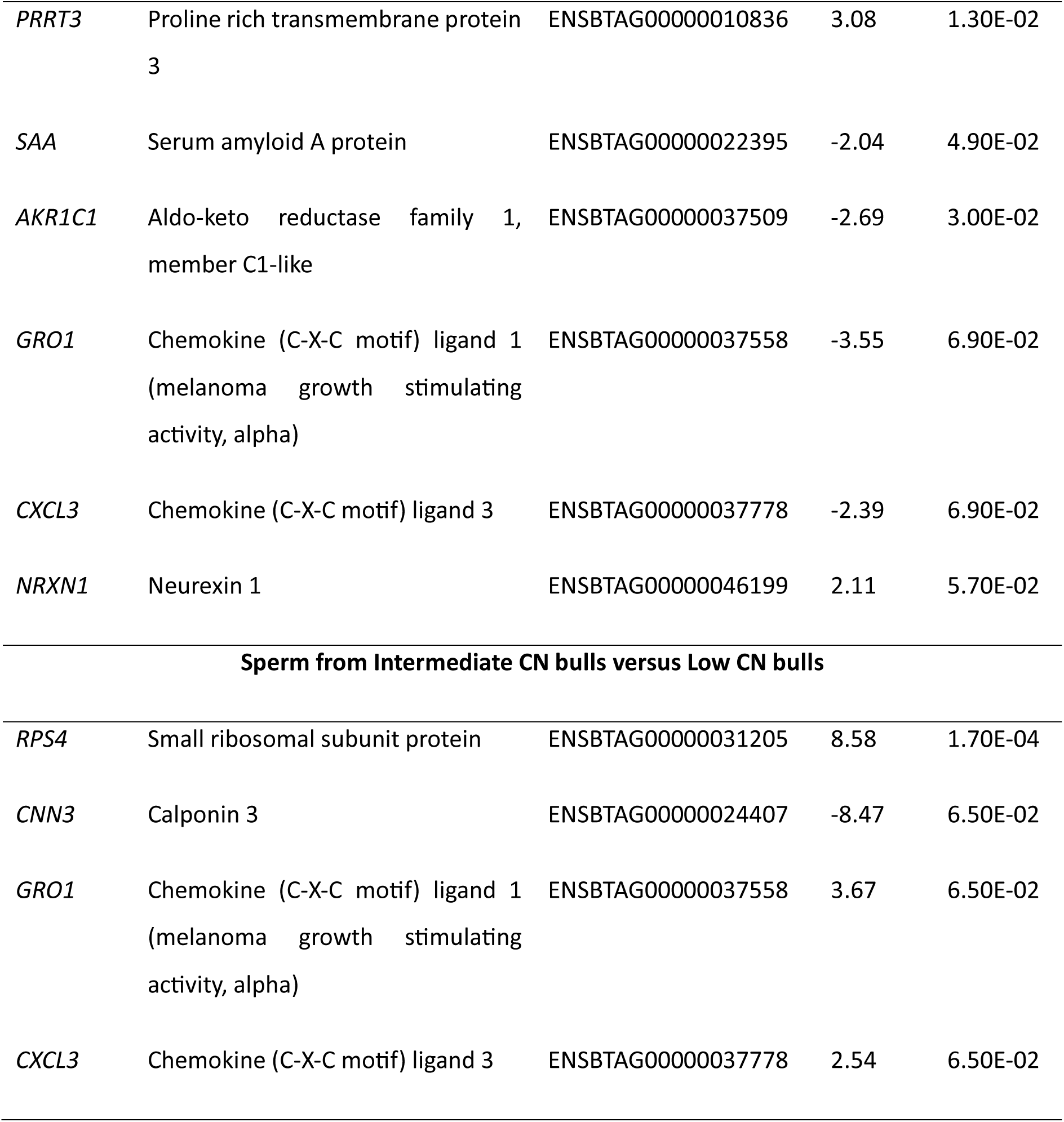
Differential gene expression analysis results comparing intermediate copy number (CN) bulls versus low CN bulls, low CN bulls versus high CN bulls and intermediate CN bulls versus high CN bull. Genes with statistically significant differences in expression (FDR<0.1) and log fold changes <-2 or >+2 were filtered for inclusion in the table.

| Gene Symbol | Description | Gene ID | Log<sub>2</sub> FC | FDR |
| --- | --- | --- | --- | --- |
| Sperm from High CN versus Low CN bulls |  |  |  |  |
| *CNN3* | Calponin 3 | ENSBTAG00000024407 | -9 | 6.70E-02 |
| *RPS3A* | Ribosomal Protein S3A | ENSBTAG00000006383 | -8.46 | 4.10E-02 |
| *CYP2C87* | Cytochrome P450, family 2, subfamily C, polypeptide 87 | ENSBTAG00000037795 | 2.6 | 9.00E-02 |
| Sperm from High CN bulls versus Intermediate CN bulls |  |  |  |  |
| *RPS3A* | Ribosomal Protein S3A | ENSBTAG00000006383 | -9.18 | 3.80E-03 |
| *BRINP3* | BMP/retinoic acid inducible neural specific 3 | ENSBTAG00000008420 | 5.53 | 8.10E-02 |
| *CDH9* | cadherin 9 | ENSBTAG00000018234 | 7.46 | 1.20E-03 |
| *RPS4* | Small ribosomal subunit protein | ENSBTAG00000031205 | -7.86 | 3.30E-03 |
| *ITLN2* | intelectin 2 | ENSBTAG00000048662 | -6.14 | 4.90E-02 |
| *GABRA4* | Gamma-aminobutyric acid type A receptor subunit alpha4 | ENSBTAG00000016645 | 4.78 | 6.30E-02 |
| *PRRT3* | Proline rich transmembrane protein<br>3 | ENSBTAG00000010836 | 3.08 | 1.30E-02 |
| *SAA* | Serum amyloid A protein | ENSBTAG00000022395 | -2.04 | 4.90E-02 |
| *AKR1C1* | Aldo-keto reductase family 1,<br>member C1-like | ENSBTAG00000037509 | -2.69 | 3.00E-02 |
| *GRO1* | Chemokine (C-X-C motif) ligand 1<br>(melanoma growth stimulating<br>activity, alpha) | ENSBTAG00000037558 | -3.55 | 6.90E-02 |
| *CXCL3* | Chemokine (C-X-C motif) ligand 3 | ENSBTAG00000037778 | -2.39 | 6.90E-02 |
| *NRXN1* | Neurexin 1 | ENSBTAG00000046199 | 2.11 | 5.70E-02 |
| **Sperm from Intermediate CN bulls versus Low CN bulls** |  |  |  |  |
| *RPS4* | Small ribosomal subunit protein | ENSBTAG00000031205 | 8.58 | 1.70E-04 |
| *CNN3* | Calponin 3 | ENSBTAG00000024407 | -8.47 | 6.50E-02 |
| *GRO1* | Chemokine (C-X-C motif) ligand 1<br>(melanoma growth stimulating<br>activity, alpha) | ENSBTAG00000037558 | 3.67 | 6.50E-02 |
| *CXCL3* | Chemokine (C-X-C motif) ligand 3 | ENSBTAG00000037778 | 2.54 | 6.50E-02 |

The greatest difference in transcriptomic response was detected when comparing uterine responses to sperm from high and intermediate CN bulls (41 genes Log2-fold-change>2). There was a decrease in the expression of 12 genes, especially *RPS3A, RPS4, AKR1C1, GRO1, CXCL3*, and *ITLN2,* in the high CN group compared to the normal CN group. In contrast, among the up-regulated genes, *BRINP3, PRRT3, GABRA4*, and *CDH9* showed the largest increases in their expression levels in response to sperm in the uterine endometrium. In comparison 3, the intermediate CN group was compared with the low CN group, a total of 6 genes were differentially expressed. Among these, *CNN3* showed an almost 64-fold lower expression, while *GRO1, CXCL3,* and *RPS4* showed significantly higher expression.

Gene ontology analysis of DEG, identified in the three pairwise comparisons, indicated cell migration as a key process influenced by *DEFB103* CN of the bull (Table 5A). Kyoto Encyclopedia of Genes and Genomes (KEGG) analysis identified several potential pathways affected – of particular interest was cell adhesion processes and the *NFKBIA/GRO1/CXCL3* signalling axis (Table 5B). Low *DEFB103* CN in bulls was associated with increased expression of genes promoting uterine preparation for embryonic development and chemotaxis. Conversely, high *DEFB103* CN was associated with the high expression of genes affecting cell adhesion and proliferation, and with low expression of genes that are crucial for protein synthesis and immune response, potentially affecting these processes in the uterine environment (all details shown in Supplementary Table S1).

**Table 5A:** Gene Ontology (GO) analysis for heifers inseminated with frozen-thawed semen from low fertility bulls, categorized by different *DEFB103* genotypes. This analysis includes all differentially expressed genes (DEGs) with a corrected p-value of less than 0.01.

| GO: Biological process |  |  |  |  |
| --- | --- | --- | --- | --- |
| Category | GO ID | Gene | Definition | P-value |
| Neural crest cell migration | GO:0001755 | *SOX8/TWIST1/S*<br>*EMA3C/SEMA3*<br>*A* | The characteristic movement of cells from the dorsal ridge of the neural tube to a variety of locations in a vertebrate embryo | 0.8E-02 |
| Mesenchymal cell migration | GO:0090497 | *SOX8/TWIST1/S*<br>*EMA3C/SEMA3*<br>*A* | The orderly movement of a mesenchymal cell from one site to another, often during the development of a multicellular organism | 0.6E-02 |

**Table 5B:** Analysis of the top 5 significant KEGG pathways for heifers inseminated with frozen-thawed semen from low fertility bulls, categorized by different *DEFB103* genotypes. This analysis includes all differentially expressed genes (DEGs) with a corrected p-value of less than 0.01.

| KEGG ID | Category | Description | Gene in pathway | P-value |
| --- | --- | --- | --- | --- |
| bta04514 | Signalling molecules and interaction | Cell adhesion molecules | *CADM3/LRRC4B/NRXN1/BOLA* | 7.20E-04 |
| bta04360 | Development and regeneration | Axon guidance | *ROBO1/ROBO2/SEMA3A/SEMA3C* | 1.02E-03 |
| bta04657 | Immune system | IL-17 signalling pathway | *NFKBIA/GRO1/CXCL3* | 1.60E-03 |
| bta04064 | Signal transduction | NF-kappa B signalling pathway | *NFKBIA/GRO1/CXCL3* | 2.60E-03 |
| bta04668 | Signal transduction | TNF signalling pathway | *NFKBIA/GRO1/CXCL3* | 3.90E-03 |

## Discussion

In this study, we investigated the effect of bull *DEFB103* CNV on sperm kinematic parameters, functional parameters and uterine transcriptomic response to sperm. This builds on previous work showing that *DEFB103* is expressed in the caput epididymis, extensive multiallelic CNV negatively correlates with its expression level in the testis and it is upregulated at sexual maturity [21]. In analysis of sperm kinematic parameters, we observed a significant negative correlation between the CN of *DEFB103* and both progressive and overall motility in bulls with HF and LF phenotypes, potentially explaining a proportion of the variation in fertility between these groups. This finding is consistent with the finding that higher *DEFB103* CN negatively correlates with gene expression in the caput epididymis [21]; consequently, reduced expression of *DEFB103* protein may impair sperm motility and function, which may explain the lower progressive and overall motility observed in bulls with higher *DEFB103* CN. Furthermore, while the CNV of *DEFB103* had a statistically significant effect on two critical sperm kinematic parameters, VSL and VAP, in bulls with LF, the differences were subtle. The observed diversity suggests that the disparities in sperm kinematic parameters cannot be attributed solely to *DEFB103* CN, highlighting the need for a multifactorial approach in analyzing bull fertility. It is possible that an additive effect across the β-defensin gene locus could have a more definitive effect.

Upon entering the female reproductive tract, sperm undergo crucial membrane transformations that are essential for fertilization [37]. Stiavnicka et al., (2023) showed that sperm from LF bulls had a reduced ability to increase membrane fluidity under *in-vitro* capacitating conditions compared to HF bulls. Our study suggests that a high CN of the *DEFB103* gene in the LF bulls may increase membrane fluidity, especially in non-capacitating conditions, which may negatively affect sperm function and contribute to the fertility differences observed among LF bulls. Additionally, research has shown differences in the ability of sperm to bind to the oviductal epithelium between HF and LF bulls [7], with β-defensins playing a critical role in this process [16,36]. Considering that high *DEFB103* CN may impair sperm function, our finding that sperm with lower *DEFB103* CN exhibit a greater ability to bind to the oviductal epithelium suggests that lower *DEFB103* CN might be associated with improved fertility outcomes by enhancing sperm binding capabilities. The increased membrane fluidity observed in low CN sperm under capacitating conditions may facilitate stronger binding to the oviductal epithelium, further supporting their role in successful fertilization.

After demonstrating a clear effect of *DEFB103* indirectly on sperm function, we sought to determine whether the uterine response varied depending on the sperm from bulls with different *DEFB103* CN. Overall, transcriptomic differences were modest, but *DEFB103* CN had significant effects on expression of genes with roles in sperm migration, immune signalling, and chemotaxis pathways. An almost 500-fold increase in the expression level of *CNN3* was detected in the Low CN group compared to others. *CNN3* regulates epithelial contractile activity by localizing to actin filaments, and interestingly, studies have shown that this is essential for embryonic development and viability in mice [32]. More importantly, *CNN3*, which showed increased expression in the low *DEFB103* CN group, has been reported to be involved in the cytoskeletal rearrangement required for trophoblastic fusion in the human BeWo cell line, a critical process for embryo implantation and interaction with the decidualized maternal uterus [38], suggesting potential novel regulatory mechanisms of defensins in semen priming the uterus for embryo implantation. *RPS4*, ribosomal protein, along with two chemokines, *GRO1* and *CXCL3*, were significantly upregulated in response to sperm from intermediate CN bulls compared to sperm from the low CN group. *RPS4* plays a role in spermatogenesis [39], suggesting that significant changes in intercellular communication, cell cycle regulation, and protein synthesis occur. *GRO1* has previously been shown to have high expression levels in decidual endometrial stromal cells in response to products secreted by trophoblasts in humans [40]. *CXCL3*, has been shown to play a role in the migration, invasion, proliferation, and tube formation of trophoblast cells [41]. Taken together, these findings suggest that *DEFB103* may contribute to chemotaxis during sperm motility and settlement within the uterus. In contrast, *ITLN2* expression was nearly 64-fold lower in the high CN group compared to the intermediate CN group. *ITLN2*, which plays a role in the innate immune response [42], may influence immune responses or tissue repair processes in the uterine environment due to its reduced expression in sperm from high CN bull. There was an increase in *CDH9* and *GABRA4* expression in sperm from high CN bulls versus intermediate CN bull group. Since *CDH9* plays a role in cell-cell adhesion [43], its increased expression in sperm may increase the interaction between sperm and the uterine epithelium. While this could potentially improve sperm retention and survival in the female reproductive tract; increased expression level of *GABRA4* may negatively affect the preparation of the uterine endometrium for implantation. It has previously been reported that *GABA* and synthetic *GABA* receptor ligands in mice can negatively affect preimplantation embryos through their receptors [44].

Taken together, the findings of this study suggest that *DEFB103* CNV exerts a significant impact on bull fertility by affecting various sperm functional parameters and eliciting distinct transcriptomic responses in the uterus post-insemination. Lower *DEFB103* CN, associated with higher *DEFB103* expression levels, likely enhances sperm motility and oviductal epithelium binding capacity by increasing protein availability and affecting charge-mediated progression to the site of fertilisation. Additionally, DEFB103 CN may alter the uterine immune response to sperm from bulls with low-CN, and therefore affecting early pregnancy establishment. These findings illustrate the impact of *DEFB103* CNV on multiple aspects of bovine reproduction and suggests that further dissection of the roles of β-defensins in cattle fertility is now warranted.

## Acknowledgements

We are grateful to Dr Miriama Stiavnicka for providing CASA and Flow cytometer data for the HF and LF phenotype bulls. We thanks to Kaitlyn Weldon and Nathan Johnston from University of Limerick for help to collect the female reproductive tract samples from the commercial abattoir. This research used the ALICE High Performance Computing facility at the University of Leicester.

## Conflict of Interest

The authors declare that they have no conflicts of interest related to the work reported in this article.

## Authorship statement

Ozge Sidekli: Conceptualisation, methodology, investigation, writing – original draft, visualisation, formal analysis, data curation, funding acquisition. Edward Hollox: Conceptualisation, supervision, writing - original draft. Sean Fair: Conceptualisation, funding acquisition, resources, supervision, writing – review and editing. Kieran Meade: Conceptualisation, funding acquisition, supervision, project administration, writing - original draft.

## Data availability

All RNA-seq data generated for this study have been deposited in the ArrayExpress database under the project accession number E-MTAB-14398.

---

**Supplementary figure S1.**
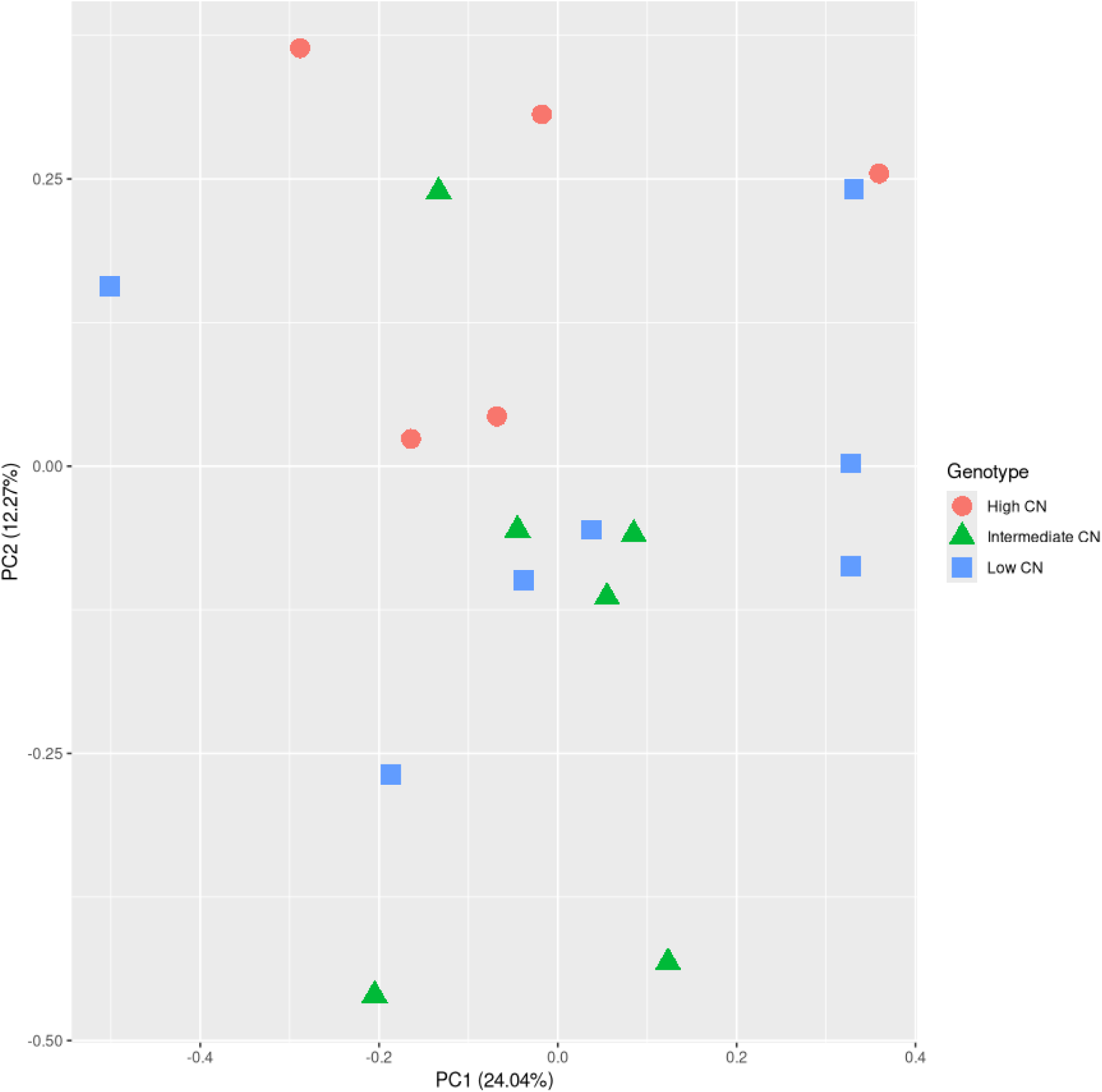
The Principal Component Analysis (PCA) plot showing the distribution of RNA-seq samples, where colours and shape indicates the three different genotypes, namely, high CN (5 heifers), intermediate CN (6 heifers) and low CN (7 heifers).

**Supplementary file S1**

## Notes

### Competing Interest Statement

The authors have declared no competing interest.

